# PET imaging for non-invasive monitoring of ^89^Zr-Talidox delivery to the brain following focused ultrasound-mediated blood-brain barrier opening

**DOI:** 10.1101/2025.06.16.659954

**Authors:** Aishwarya Mishra, Chris Payne, Amaia Carrascal-Miniño, Stefan Halbherr, Antonios N. Pouliopoulos, Rafael T. M. de Rosales

## Abstract

The blood-brain barrier (BBB) significantly hinders the treatment of central nervous system (CNS) disorders and brain tumors with intact BBB by restricting the entry of most therapeutic agents, including small-molecule drugs and particularly larger macromolecules. Liposomal formulations, such as PEGylated liposomes with long blood half-lives, high drug-carrying capacity, and reduced off-site toxicity, can be useful for brain drug delivery, but their large size often limits BBB penetration. A novel liposomal doxorubicin formulation, Talidox®, with a smaller size (∼36 nm), increased blood circulation half-life, and better stability than previous clinical formulations, can be a suitable choice for brain delivery. This study investigated Talidox® delivery to the brain through focused ultrasound (FUS)-mediated BBB transient opening. Radiolabelling of Talidox® via intraliposomal ^89^Zr enabled Positron Emission Tomography (PET) imaging for whole-body non-invasive, real-time monitoring of biodistribution and pharmacokinetics. Following FUS-mediated BBB opening in mice, PET imaging revealed a significant increase in brain uptake compared to non-FUS controls, achieving a 14-fold higher accumulation. Additional validation using passive acoustic detection, microscopy, autoradiography, and cryo-fluorescence tomography demonstrated successful brain distribution that correlated with PET imaging results. These findings underscore the potential of combining Talidox® with FUS for effective, non-invasive drug delivery to the brain and highlight the advantages of PET imaging as a modality for non-invasive, longitudinal quantification of drug delivery to the brain.

## Introduction

The blood-brain barrier (BBB) hinders advancement in delivering therapeutics to specific brain areas. Many central nervous system (CNS) pathological conditions and brain tumors with intact BBB, such as glioblastoma (GBM), remain untreatable or partially treatable.^1,2^ The BBB blocks more than 98% of small-molecule drugs and all larger therapeutic molecules, rendering them suboptimal for treating CNS ailments.^3^ Moreover, drug delivery to the brain is affected by further issues, including non-spatial targeting, non-specific toxicity, subtherapeutic drug-dose accumulation, and metabolic degradation.^4,5^

Drug delivery approaches for encapsulating small molecule therapeutics offer great promise for targeted brain treatment. Liposomes are widely used for drug delivery due to their ability to target specific areas while minimizing systemic toxicity.^6^ Liposomal nanomedicines can carry hydrophilic and hydrophobic drugs at higher concentrations, can be modified to target specific sites, and, when PEGylated, increase blood circulation half-lives.^7,8^ Additionally, they offer theranostic capabilities^9^ and targeted release mechanisms.^10^ Although most research has focused on oncology, liposomes also show potential for delivering small-molecule drugs to treat neurodegenerative diseases.^11^ However, their large size typically prevents them from crossing the BBB. Efforts to optimize size for brain delivery and further minimize the toxicity of PEGylated liposomal doxorubicin have led to the development of TLD-1 (Talidox®), a novel formulation with smaller, uniform, more stable liposomes (36 nm average diameter compared to 70 nm for Caelyx®), potentially improving its safety profile.^12,13^

Optimizing liposomes for BBB penetration has significant implications for treating CNS diseases. Strategies to enhance liposomal formulations for BBB crossing include cationization,^14,15^ active targeting,^16,17^ stimuli-responsive,^18^ and multifunctional liposomes^19^ but with limited success.^20^ Various invasive and non-invasive methods have been investigated to overcome the BBB.^21,22^ Invasive methods, such as direct brain injections, can cause neurological damage, infection, and bleeding.^23,24^ Osmotic^25^ and chemical^26,27^ disruption methods have also been tested but suffer from drawbacks and have failed in clinical trials. Modifying drugs to utilize endogenous transport mechanisms for BBB crossing has shown limited success in achieving therapeutically relevant brain concentrations.^20^

A non-invasive approach using focused ultrasound (FUS) and microbubbles (lipid-shelled gas particles with a heavy gas core and a diameter of 1-10 µm) has shown promise for targeted delivery to the brain.^28^ This technique involves injecting microbubbles and therapeutic agents into the bloodstream and then applying ultrasound pulses to the targeted brain region. The oscillation of microbubbles facilitates drug delivery across the BBB.^29,30^ Previous work in this area has demonstrated that liposomes can be delivered into the brain using this focused ultrasound technique.^31–47^ The cargo of these liposomes included genes,^35,38,39^ chemotherapeutic drugs,^32,36,37,42,43^ and imaging agents,^33,37,40,41,45,46^ with diameters ranging between 55 and 200 nm. The above studies have shown that the extent of delivery decreases with larger liposome sizes, consistent with the BBB’s protective role against administered agents.

Interestingly, most of the studies discussed above demonstrate that focused ultrasound-mediated BBB opening facilitates the delivery of various liposomal formulations. However, these studies primarily use optical imaging or MRI to assess the extent of drug delivery, which limits our ability to understand the whole-body pharmacokinetics of the drug delivery system and accurately quantify drug delivery, particularly at the human-body scale.^48^ Positron Emission Tomography (PET) imaging can improve our ability to measure drug distribution, uptake, and pharmacokinetics of drugs within the brain on account of its sensitive and quantitative nature.^49^ PET employs radiolabelling of drugs and drug-delivery vehicles followed by their administration *in vivo* as radiopharmaceuticals and quantifying the radiopharmaceutical delivered to the tissue during the scan.^50^ Moreover, PET is sensitive in its measurements of small changes in tissue accumulation and regional cerebral metabolism clinically.^51^ Consequently, PET has been used to monitor transport of various drugs across the BBB, including small molecules,^52^ nanoparticles,^53,54^ viral vectors,^55^ and antibodies.^56^

We hypothesized that Talidox®, a novel liposomal formulation with an average size of 36 nm and improved safety profile,^13^ can be delivered to the brain using focused ultrasound at therapeutic concentrations. The radiolabelling of Talidox® with ^89^Zr facilitated the use of PET imaging to quantify and monitor whole-body biodistribution and pharmacokinetics of liposomal delivery through FUS-mediated BBB opening for the first time. Brain delivery of doxorubicin across the BBB was visualized using fluorescence imaging in both FUS and non-FUS control subjects. The effect of Talidox’s long blood half-life (t_1/2_ = 95 h) on accumulation through the BBB opening, which remains open for at least 24 h, was also examined. The non-invasive, longitudinal and quantitative nature of PET imaging used here to determine the extent of BBB opening and liposomal dose accumulation within the brain in this study will pave the way for future therapeutic studies using combination of Talidox® and FUS.

## Methods and Materials

### Materials

Deionized water was obtained from a PURELAB® Chorus 1 Complete instrument (Veolia Water Systems LTD, UK) with 18.2 MΩ cm resistance and was used throughout this study. Talidox® was provided by InnoMedica Holding AG (Switzerland). ^89^Zr was purchased from PerkinElmer as [^89^Zr]Zr-oxalate in 1.0 M oxalic acid (BV Cyclotron, VU, Amsterdam, the Netherlands). Radioactivity in samples was measured using a CRC-25R dose calibrator (Capintec). All other chemicals and reagents were commercially available, of analytical grade, and used without further purification.

### Synthesis of [^89^Zr]Zr(oxinate)_4_ via kit method

^89^Zr (4-7 MBq, 6 µL) in 1 M oxalic acid (PerkinElmer) in a screw cap glass vial, followed by addition of 54 μL of aqueous buffered oxine kit (8-hydroxyquinoline, 8HQ) solution containing 0.5 mg/mL 8HQ and 1 mg/mL Tween-80, and 1 M HEPES at pH 7.8. pH was measured at ∼7.2-7.5, and the radiolabelling mixture was incubated at RT for 10 min.^53^

[^89^Zr]Zr(oxinate)_4_ complex formation was confirmed by iTLC; stationary phase = Whatman No 1 filter paper (Cytiva, USA) and mobile phase = 100% ethyl acetate. The chromatograms were analyzed on Scan-RAM interfaced with a PET probe and processed using Laura ITLC software (Lablogic Systems Ltd, United Kingdom). Regions of interest (ROI) were drawn from R_f_ = 0.5-1 to quantify the area under the curve. A radiochemical purity (RCP) of [^89^Zr]Zr-oxine above 85 % was deemed acceptable to proceed to radiolabelling of the liposomes.

### Radiolabelling of Talidox

20 µL of [^89^Zr]Zr(oxinate)_4_ complex was added to 200 µL of Talidox® in 280 µL of phosphate-buffered saline (PBS) and incubated at 50°C for 30 min under continuous shaking. The mixture was then purified using a PD Minitrap™ G-25 size exclusion column (Cytiva, USA) following the manufacturer’s gravity protocol. In brief, 500 µL of liposomal radiolabelling mixture was added to a previously PBS equilibrated column, and the purified liposomes were eluted with 750 µL of PBS, discarding the first 200 µL, and purified radiolabelled liposomes were obtained in the last 550 µL. Radiochemical yield (RCY) was calculated as shown below (CPM = counts per minute):

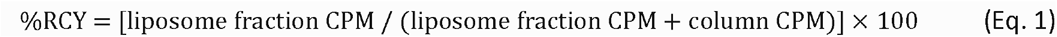

For *in vivo* experiments, purified radiolabelled liposomes (550 µL) were concentrated to their original volume (200 µL) using centrifugal size exclusion spin filtration (Amicon® Ultra 0.5 mL 30 K filters (Millipore, Merck, Germany)).

### Quality Control of [^89^ Zr]Zr-Talidox

Quality control of the liposomes involved dynamic light scattering (DLS) hydrodynamic size, polydispersity index (PDI) and zeta-potential measurements, performed with Zetasizer Nano ZS (Malvern, United Kingdom). All the above measurements were performed pre- and post-labelling of liposomes.

The radiochemical purity (RCP) of the [^89^Zr]Zr-Talidox was determined using a centrifugal size exclusion spin filtration (Amicon® Ultra 0.5 mL 30 K filters (Millipore, Merck, Germany)). Any non-specific bound radioactivity was collected in the filtrate, and liposome-bound radioactivity was retained in the filter. The radiolabelled liposomes were only used for further experiments if the RCP was higher than 95%.

#### Serum stability of [^89^Zr]Zr-Talidox

[^89^Zr]Zr-Talidox (200 µL) was incubated in human serum (200 µL) in a 1:1 ratio for up to 72 h. Aliquots of the test sample were taken at different time points for stability study and applied to SEC HPLC (ÄKTA purifier (GE, Sweden) equipped with a Superose™ 6 Increase 10/300 GL (Cytiva, USA) column running with 0.5 mL/min in PBS) at 0, 24 h, 48 h and 72 h. Thirty 1 mL fractions were eluted in PBS. UV samples were measured at 280 nm to determine fractions containing liposomes. The radioactivity signal of samples was recorded on a gamma counter (LKB Wallac 1282 Compugamma S) to determine the radioactivity associated with liposomes and serum. The serum stability % was determined using the equation below (CPM = counts per minute):

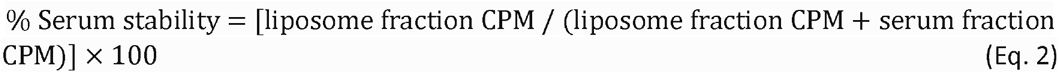

### Animals and study design

All animal experiments were ethically reviewed by the Animal Welfare & Ethical Review Board at King’s College London and carried out in accordance with the Animals (Scientific Procedures) Act 1986 (ASPA) UK Home Office regulations governing animal experimentation. The animal studies were performed under project license PPL PBBA9A243. All *in vivo* experiments were conducted on healthy female BALB/c mice (8–9 weeks old) obtained from Charles River UK Ltd.

Ten mice were used to compare the dose and distribution of liposomal delivery in animals treated with focused ultrasound (n = 4) and untreated animals (n = 6). The brain treatment’s acoustic pressure and other ultrasonic parameters were chosen based on previous optimization experiments using the same pre-clinical FUS system.^47^

Control untreated animals (n = 6) were divided into two groups: *(a)* administration of radiolabelled liposomes ([^89^Zr]Zr-Talidox liposomes: 80-100 µL, 1 mg/mL Dox, 0.6-1 MBq) without microbubbles (n=3); and *(b)* administration of microbubbles with radiolabelled liposomes ([^89^Zr]Zr-Talidox liposomes: 80-100 µL, 1 mg/mL Dox, 0.6-1 MBq) (n=3).

Treated animals (n = 4) were administered with microbubbles during ultrasound treatment followed by the administration of radiolabelled liposomes ([^89^Zr]Zr-Talidox liposome: 80-100 µL, 1 mg/mL Dox, (0.6-1 MBq)).

### Microbubbles

BR1 microbubbles (Bracco S.p.A., Milan, Italy) were used to deliver liposomes into the brain (concentration: 5 µL/g (1.5 × 10^9^ MB/kg) of body mass, volume: 100 µL, mean diameter: 2.5 µm, vial concentration: 3x10^8^ /mL). BR1 is the research-grade analogue of SonoVue®, which is routinely used in contrast-enhanced ultrasound imaging in the clinic. A fresh vial of microbubbles was used on each day of experiments and used within six hours of first use, following the manufacturer’s instructions.

### Ultrasound setup and experimental conditions

FUS experiments were conducted in the same experimental setup as described in detail previously.^47^ Briefly, a 0.5-MHz single-element FUS transducer (Part No. H-204; Sonic Concepts, Bothell, WA, USA) was driven by a waveform generator (33500B series; Agilent Technologies, Santa Clara, CA, USA) through a radiofrequency power amplifier (A075 RF Power Amplifier, 300 kHz to 35 MHz, 75 W; E&I, Rochester, NY, USA). Acoustic emissions were captured with a 7.5-MHz single-element passive cavitation detector (Part No. U8423042, V320-SU, diameter: 12.7 mm; Evident Europe GmbH – UK Branch, Stansted, UK) which was inserted and co-aligned with FUS transducer. A high-pass filter was used to filter out the fundamental and the second harmonic reflections (Part No. ZFHP-1R2-S+, cut-off frequency 1.2 MHz; Mini Circuits, Brooklyn, NY, USA). Recorded signals were amplified by 20-dB with a pulser-receiver (VK-71000, Gampt mbH, Merseburg, Germany) and then recorded using a Picoscope (5244D, Picotech, UK). Segments of 187,500 time points were captured at a sampling frequency of 125 MSa/s.

Anesthesia was induced and maintained with inhalable isoflurane mixed with oxygen (∼4% induction, 1.5-2% maintenance) delivery through a digital vaporizer (Somnosuite, Kent Scientific, Torrington, CT, USA). Mice were fixed in a stereotactic frame for accurate targeting. Head fur was removed with clippers and depilatory cream applied for 10-20 seconds. Ultrasound gel was applied to the mouse head, and a 3D-printed square container with a transparent base was filled with degassed water and placed onto the mouse head to visualize the sutures of the skull for targeting purposes. An ultrasound transducer mounted with a cone filled with continuously degassed distilled water and enclosed with a parafilm membrane was lowered into the water bath. For targeting purposes, a 1-mm-thick metal cross was placed at the bottom of the water bath and in alignment with the lambdoid and sagittal sutures of the skull. Using a metallic grid method previously described,^58^ the caudate area was targeted (-2 mm lateral, +2 mm ventral from lamboid suture). Caudate was targeted as it is a common tumor location. A control sonication was performed before microbubble injection to acquire a baseline signal, which was subsequently subtracted from the microbubble signal in the frequency domain. BR1 microbubbles were administered intravenously via a catheter inserted into the tail vein at a 1:4 dilution with a total bubble dose of 2 × 10^7^ per injection. Mice were then treated using the following sonication parameters: peak-negative pressure: 320 ± 30 kPa, centre frequency: 0.5 MHz, pulse length = 500, cycles = 1 ms, Pulse repetition frequency = 5 Hz, total number of pulses = 600. The opposite right side acted as a control (no ultrasound treatment). Following sonication, [^89^Zr]Zr-Talidox liposomes were administered using the same catheter.

### Passive Cavitation Analysis

Acoustic cavitation emissions were processed using Matlab® (Mathworks, MA, USA), similarly to previous studies.^59,60^ Briefly, the energy emitted from exposed microbubbles during a therapeutic ultrasound pulse was estimated from the time domain signal through:

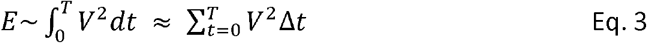

where V was the voltage at each time point in volts, and the sampling period was 1.25 x 10^-8^ s. The circuit energy detected was assumed to be proportional to the acoustic energy emitted by microbubbles exposed to therapeutic ultrasound. Control sonications before microbubble administration were used to estimate the baseline acoustic signal due to reflections. The average energy of control sonications was subtracted from the microbubble signal for each pulse.

Frequency analysis was used to identify the dominant cavitation mode for each treatment.

Three spectral areas from the Fast Fourier Transform (harmonic region, ultraharmonic region, and broadband regions) were filtered and identified in the frequency domain:

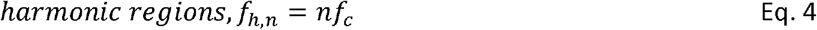

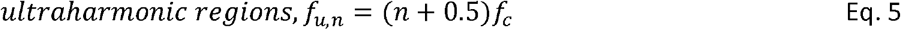

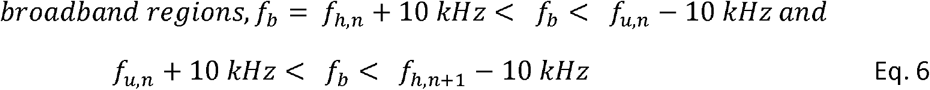

where f_c_ is the centre frequency of the FUS transducer and n is the harmonic/ultraharmonic number. The fundamental and second harmonic were filtered out using a 1.2-MHz high pass filter. Cavitation doses were calculated based on the root-mean-square voltage detected in the retrospective areas.^61^ The stable harmonic (SCD_h_), ultraharmonic (SCD_h_ ), and inertial (ICD_u_) cavitation doses were defined as:

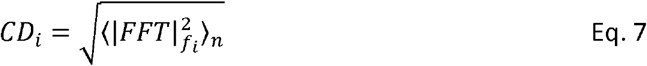

where *i* changes for harmonic (n = 3 to 10), ultraharmonic, and broadband (inertial) frequency domains.

### PET imaging and Biodistribution

Post administration of the radiolabelled liposomes ([^89^Zr]Zr-Talidox) via a tail vein intravenous catheter, mice were scanned in a preclinical NanoScan® PET/CT imaging system (1:5 coincidence mode; 5-ns coincidence time window; Mediso Medical Imaging Systems, Budapest, Hungary) using 4-bed hotel multi animal imaging setup at the following time points post-administration of [^89^Zr]Zr-Talidox: t = 1, 4, 24, 48 and 72 h for a scan time duration of 1 h. The mice were kept anesthetized inside the scanner at 1.5–2.0% vaporized isoflurane mixed with oxygen. PET/CT images were reconstructed using Tera-Tomo 3D reconstruction (400–600 keV energy window, 1–3 coincidence mode, 4 iterations, and subsets) at a voxel size of (0.4 × 0.4 × 0.4) mm^3^ and corrected for positron range, attenuation, scatter, and decay.

The reconstructed images were analyzed using InterView™ FUSION processing software (Mediso) and VivoQuant (Perceptive (formerly Invicro)) for image analysis. The four animal PET/CT images acquired were segmented into individual animal PET/CT images via Interview Fusion. All individual images thus obtained were separately saved as DICOM files with their respective injected dose and animal weight before performing ROI analysis.

Region of interest analysis was performed using VivoQuant V2.5 Patch 3 Software with a 3D Brain Atlas plugin module. The whole-body PET images were cropped to obtain the head PET/CT image, followed by registration of a pre-defined brain atlas (containing 28 predefined brain regions) to the head PET/CT image, providing region-specific volume and uptake quantification. The head PET/CT images were presented as PNG files with or without the skull in different orientations to highlight both treated and untreated contralateral regions.

Post scanning at t=72 h, the mice were culled by cardiac puncture under anesthetic overdose (>5% isoflurane/oxygen) followed by perfusion with PBS and 4% PFA to allow for removal of any blood circulating liposomes as well as the preservation of the brain tissue structure. Multiple organs were collected, weighed, and measured for radioactivity using a Wallac gamma counter. Serial standard dilution of the injected radiotracer was measured alongside the collected organs to calculate the percentage injected dose (% IA g^−1^).

### Autoradiography

Isolated perfused brains, either treated or untreated, were placed in 4% PFA for 24 h and transferred to 30% sucrose for cryoprotection. For autoradiography, these cryoprotected brains were coated with OCT, placed into an OCT-filled mold, and placed in dry ice. The mold containing the brain was mounted on the cryostat, and 3 x 20 µm sections per Superfrost slide were obtained along the axial plane from the top to the bottom of the brain.

The treated and untreated brain sections were exposed to a phosphor storage screen. The Amersham TYPHOON scanner detected the developed phosphor storage screen with accumulated radiation energy, creating a digital image of the radiated areas of the storage phosphor screen. The obtained digital images were analyzed using the software ImageQuantTL 10.0 -261 (Cytiva, USA) or ImageJ. A 3D reconstruction of images from a single brain was constructed using sequential sections using a 3D plugin in ImageJ.

### Microscopy

The sections of treated and untreated brains used for autoradiography were further stained with DAPI and acquired using fluorescence microscopy. The slices were imaged with an EVOS™ M5000 Imaging System (Thermo Fisher Scientific, USA) equipped with DAPI (Ex: 357/44 nm; Em: 447/60 nm) and RFP (Ex: 542/20 nm; Em: 593/40 nm). The doxorubicin incorporated in the liposome formulation was detected in the RFP channel. The stained nuclei were detected in the DAPI channel. The contralateral side of the brain in treated brains served as an intrasubject control. Non-sonicated untreated brains were also imaged as a control. The images acquired in different channels were also overlaid to obtain a composite image. Images were processed using either ImageJ or Fiji.

### Cryo-fluorescence Tomography

Isolated perfused brains (1 x sonicated treated brain, 1 x non-sonicated untreated brain, 1 x control brain) stored in H_2_O were flash-frozen for analysis using cryo-fluorescence tomography (CFT) (Xerra, EMIT) to validate the location of liposomes by visualizing the fluorescence signal of doxorubicin. Excised brains were frozen over dry ice and then embedded top-down in the transverse plane in OCT in the smallest predefined mold available. White light and fluorescence images (excitation 470 nm, emission 620 nm, 1500 millisecond exposure) were acquired with 30 µm in-plane resolution and 50 µm slice thickness. Images were combined into stacks using ImageJ (Fiji) and then visualized in the 3D plugin to create maximum-intensity projections.

### Statistics

Descriptive data are presented as mean ± standard deviation unless otherwise mentioned. Results were considered statistically significant at p < 0.05, without correction for multiple comparisons. A comparison of groups, analysis, and graph generation was performed using GraphPad Prism version 10 (GraphPad Software 10.4).

## Results and discussions

### Talidox® liposomes can be radiolabelled with high stability

The positron emitter radionuclide ^89^Zr (*t*_1/2_ = 78.4 h) was selected over other short-lived isotopes such as ^68^Ga (*t*_1/2_ = 67.7 min) and ^18^F (*t*_1/2_ = 109.8 min) because its half-life is similar to the half-life of circulating liposomes such as Talidox® (median half-life of 95 h).^13^ Moreover, ^89^Zr was also chosen over other long-lived radionuclides such as ^64^Cu (*t*_1/2_ = 12.7 h) because free ^64^Cu shows similar biodistribution and excretion routes as liposomes and increased brain accumulation, whereas free ^89^Zr accumulation is not observed in the liver or brain.^62^ Thus, ^89^Zr simplifies the interpretation of PET imaging studies tracking liposomal nanomedicines, especially at later time points when the release of free radiometal is expected.^63^

The radiolabelling of Talidox® with ^89^Zr was a two-step process: synthesis of [^89^Zr]Zr-oxine via a single step kit-based method^57^ followed by co-incubation with Talidox®. This radiolabelling method is chelator-free and requires no chemical modification of the liposomal formulation^64^, thereby minimizing the impact on the physicochemical properties of the original liposomal formulation. [^89^Zr]Zr(oxinate)_4_ , a metastable, neutral, and lipophilic complex, was used to passively transport the radionuclide inside the liposomal core (Figure 1A). This metastable complex inside the core of the liposomes is further stabilized by the presence of the encapsulated drug doxorubicin. The long-term radiochemical stability of such formed radiolabelled liposomes depends on the interaction between the drug and radiometal.

**Figure 1.**
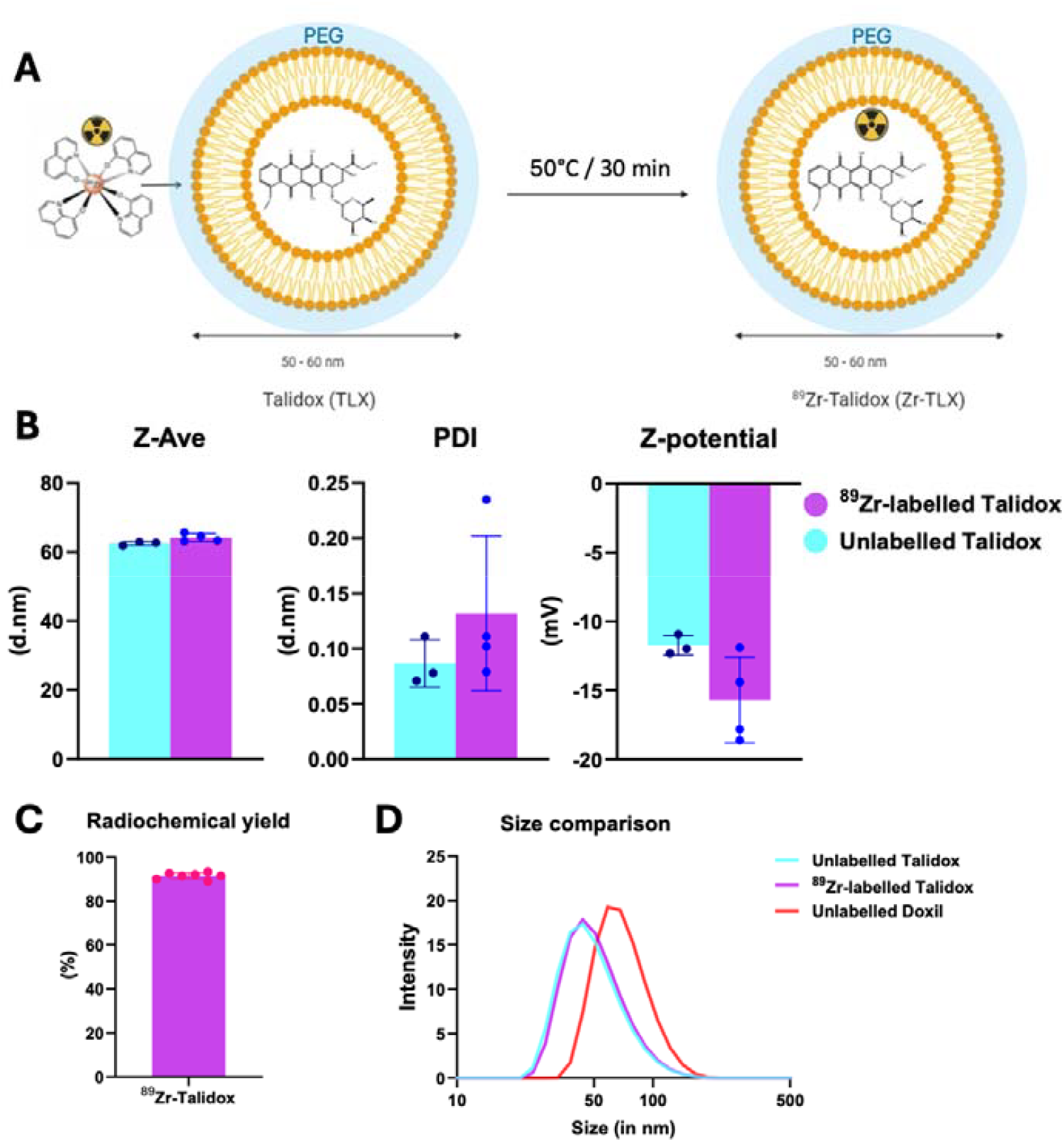
Synthesis and Characterisation of [ ^89^Zr]Zr-Talidox liposomes: **(A)** Radiolabelling scheme using kit synthesised ^89^Zr-oxine and Talidox liposomes; **(B)** Characterisation of Talidox before and after ^89^Zr radiolabelling: (i) average hydrodynamic size (Z-average) remained unchanged pre- and post-radiolabelling (p = 0.08), (ii) Polydispersity Index (PDI) of Talidox was unaffected by radiolabelling (p = 0.3), and (iii) Surface charge (Z-potential) of Talidox was also unaffected by radiolabelling (p = 0.09); **(C)** Radiochemical yield of Talidox radiolabelling reaction (n = 7); **(D)** Size comparison between the conventional formulation Doxil liposomes, smaller novel formulation Talidox liposomes, and ^89^Zr-Talidox liposomes.

The radiochemical yield of the radiolabelled liposomes obtained in feasibility studies was 91 ± 2% (n=4) for Talidox® (Figure 1C). The radiochemical yield decreased with decreasing purity of the synthesized [^89^Zr]Zr-oxine complex, and therefore, only batches with higher than 85% RCP were utilized for the radiolabelling of liposomes. Tween 20, which is one of the kit components and a surfactant, could affect the structural integrity of the liposomes in high concentrations. Therefore, the total concentration of kit components in the liposome radiolabelling mixture was never allowed to reach more than 5% to avoid any impact on the structural integrity of the Talidox® liposomes.

The stability of [^89^Zr]Zr-Talidox in human serum post 72 h incubation at 37°C was 75 ± 11% for [^89^Zr]Zr-Talidox. This stability is comparable to known radiolabelling studies of liposomal doxorubicin formulations. The slight amount of unbound [^89^Zr]Zr, probably not internalized during labelling and present in the phospholipid bilayer, is transchelated by proteins in the serum and removed from the liposomes. Interestingly, such *in vitro* serum stabilities have been consistently observed in studies involving *in vivo* tracking of liposomes without impact on *in vivo* biodistributions.^64^ Moreover, free [^89^Zr]Zr is not known to accumulate in the brain. Therefore, the stability was deemed enough to progress into *in vivo* studies.

### Radiolabelling does not affect Talidox properties

The physicochemical properties of liposomal nanomedicine formulations play an essential role in their drug delivery ability. Talidox was chosen over other widely used liposomal formulation, such as Doxil, for its higher stability, drug concentration, and smaller size, which make it an ideal choice for delivery to the brain. Therefore, these physicochemical properties were considered essential to be conserved post-radiolabelling.

Radiolabelling did not affect the size (hydrodynamic diameter), surface charge (z-potential), or size distribution (polydispersity index) of Talidox (Figures 1B and 1D). The hydrodynamic diameter of Talidox was 62.5 ± 0.5 nm and 64 ± 1 nm before and after labelling, respectively (P = 0.08), the z-potential was -5.9 ± 1.1 mV and -3.5 ± 1.8 mV (P = 0.09) before and after the labelling (Figure 1B) and finally, polydispersity index was 0.08 ± 0.02 and 0.13 ± 0.07 before and after labelling with [^89^Zr]Zr (P = 0.3). The above results thereby confirmed that the properties of the Talidox are conserved after the radiolabelling as expected on account of the chelator-free and internal radiolabelling method utilized.

### PET imaging of ultrasound-mediated Talidox delivery to the brain

After the successful synthesis of [^89^Zr]Zr-Talidox and establishing their serum stability, the radiolabelled liposomes were tracked in healthy BALB/c mice to determine the baseline delivery of Talidox that can be achieved without any interventions to open the BBB. The radiolabelled liposomes were administered intravenously and imaged at different time points (Figure 2A,B) followed by *ex vivo* tissue biodistribution analysis at the end of the imaging session (Figure 2C). Talidox liposomes were observed in circulation in the vasculature and heart for up to 24 h. During later time points, the accumulation showed similar biodistribution with other liposomal formulations, with uptake observed in mononuclear phagocytic system organs such as the spleen and liver. Due to the release of small amounts of free ^89^Zr, low levels of bone uptake were observed. However, imaging confirmed the longer circulation times in blood as expected for this novel formulation of Talidox compared to earlier imaging studies performed on other liposomal formulations.^12,13^

**Figure 2.**
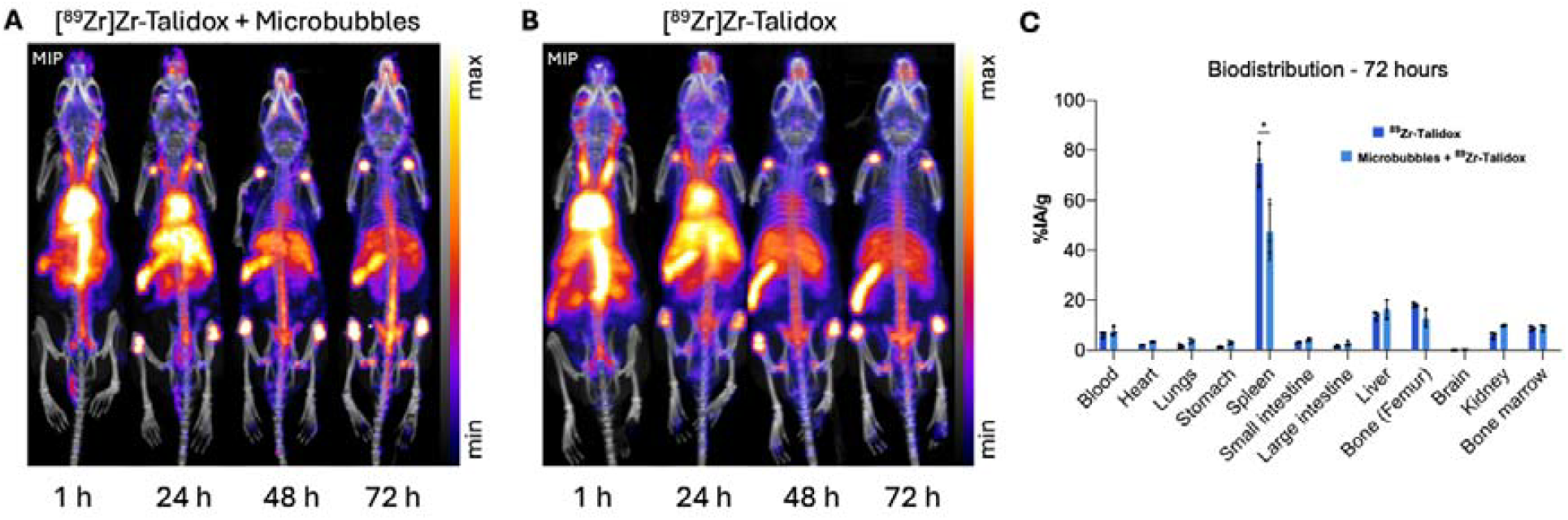
Whole body PET imaging and biodistribution of [^89^Zr]Zr-Talidox in healthy BALB/c mice: **(A**,**B)** Longitudinal PET imaging over 72 h showing biodistribution of [^89^Zr]Zr-Talidox in the presence (A) and absence (B) of microbubbles; **(C)** Biodistribution performed post-imaging showing the difference in spleen accumulation of [^89^Zr]Zr-Talidox in the presence of microbubbles (p<0.05).

PET Image-based analysis of our target organ, the brain, showed limited signal (<2% IA/g) at the earliest time point post-administration, followed by a decrease in uptake at each time point and finally, less than 1% IA/g at 72 h. Post-mortem biodistribution confirmed these results with whole brain uptake of 0.05 ± 0.02 % IA/g. The differences observed between the image quantification data and post-mortem biodistribution are due to transcardial perfusion, which allowed the removal of blood-circulating Talidox liposomes, leading to an overestimation of the dose delivered to the brain in the image quantification. Further analysis of the PET images showed that the uptake ratio observed between the left and right hemispheres of the brain was unity as expected. This quantification was particularly useful due to the contralateral control in the treated mouse brains. The control group showed that even with the improved properties of Talidox®, high delivery to the brain cannot be achieved without a BBB opening intervention.

BBB opening using focused ultrasound involves the administration of microbubbles before the application of ultrasound. Before investigating the impact of BBB opening on the delivery of liposomes to the brain, we aimed to establish the impact of the co-administration of microbubbles and [^89^Zr]Zr-Talidox liposomes on the whole-body biodistribution of [^89^Zr]Zr-Talidox. In past studies involving focused ultrasound for drug administration to the brain, the impact of microbubble administration on whole-body biodistribution of drugs has never been investigated. However, by using PET imaging, we can non-invasively and longitudinally examine the potential effects of microbubble administration on drug biodistribution. To this end, in this control imaging group, mice were administered with microbubbles before administration of [^89^Zr]Zr-Talidox.

The imaging and biodistribution results confirmed no interference between microbubbles and liposomes, as the long blood circulation properties of the liposomes were unaffected, and the observed brain uptake was also unaffected (Figure 2C). However, the post-mortem biodistribution at 72 h showed decreased uptake in the spleen (47 ± 11% IA/g) compared to the previous group (75 ± 8% IA/g). This could be attributed to the accumulation of microbubbles in the spleen, which could lead to its saturation. However, microbubbles are considered truly vascular agents. The accumulation of microbubbles in the spleen could be due to the reticuloendothelial system, but that would lead to accumulation in the liver, which was not observed. The splenic tropism of microbubbles, as seen previously in a healthy human volunteer study using microbubbles as contrast agents for ultrasound imaging,^65^ further supports our assertion that decreased splenic uptake of [^89^Zr]Zr-Talidox is due to increased spleen uptake of microbubbles. This hypothesized indirect observation of increased microbubble accumulation in the spleen and its dependence on administered microbubble concentration requires further exploration. This phenomenon of splenic tropism could potentially be exploited to block spleen uptake of cancer nanomedicines and therapeutics in the clinic.

Finally, the test group involved opening the BBB using focused ultrasound and microbubbles followed by administration of the [^89^Zr]Zr-Talidox (See Figure 3 for FUS setup, including sonication area (B) and imaging plan (C)). Focusing on the brain images showed a hot spot on the treated side of the brain, whereas no hotspots were seen on the contralateral control side. The hot spots were persistent and were observed even at 72 h (Figure 4). The contrast of these spots increased over time, showing that the signal is due to labelled liposomes that have crossed the BBB, not liposomes still in blood circulation. The PET signal decreased with increasing depth in the brain. Determining the penetration depth of the liposomes over time is useful in determining the extent of delivery in different brain areas. Interestingly, the radioactive signal within the brain appears to diffuse over time. This is expected due to the small size of Talidox liposomes (∼ 36 nm), which could allow them to diffuse through the ∼64 nm pores of the extracellular matrix of the brain.^66^

**Figure 3.**
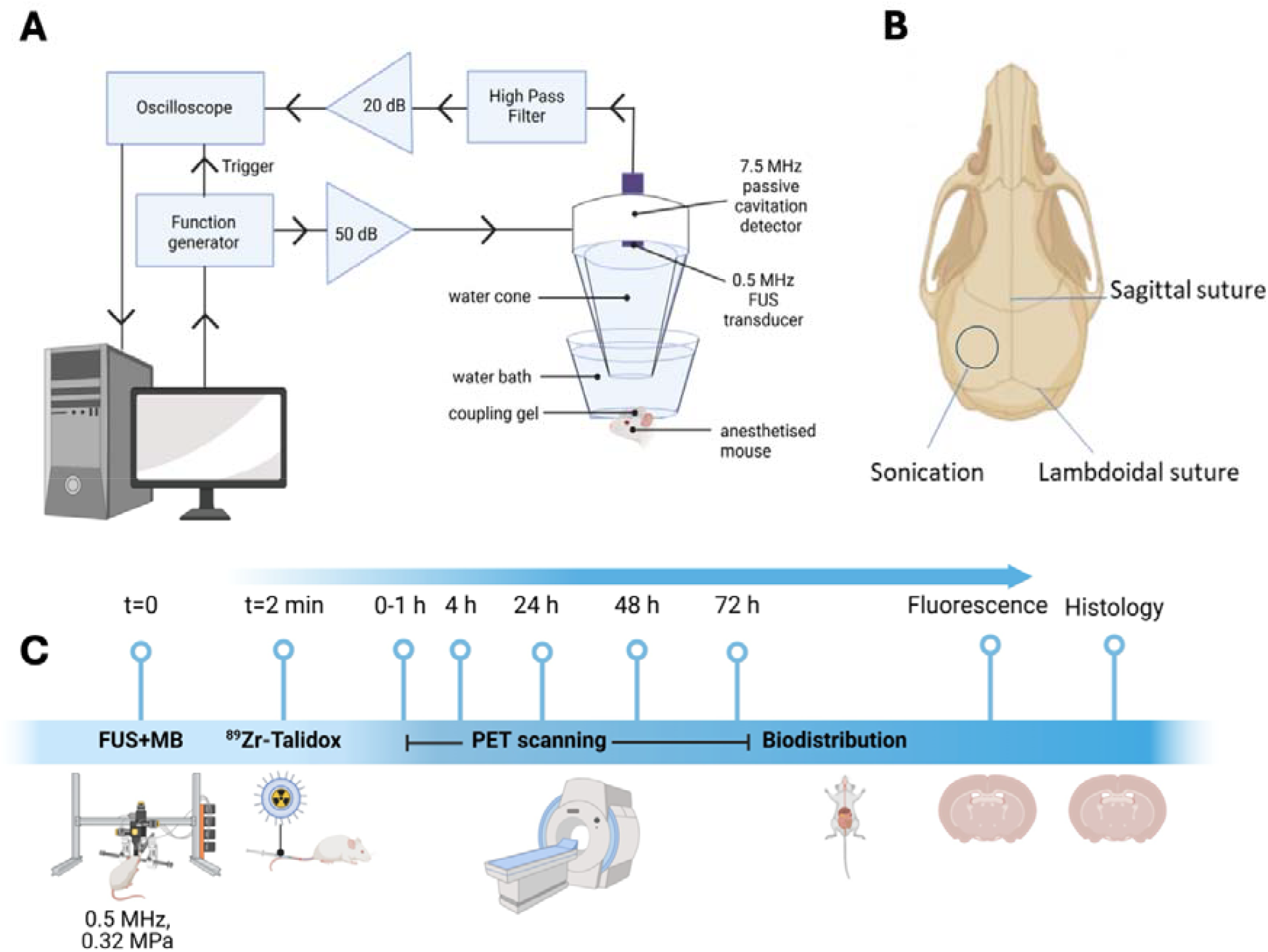
Schematics and planning of the focused ultrasound sonication experiment: (A) Focused ultrasound pulses were administered using 0.5 MHz ultrasound transducer. The ultrasound emissions from microbubbles was recorded using a 7.5 MHz passive cavitation detector; (B) The sonication was focused through the intact skull onto the left hemisphere of the brain whereas the right untreated side acted as a contralateral control; (C) Schematic plan of the FUS+PET experiments for tracking the delivery of ^89^Zr-Talidox through the FUS mediated BBB opening involving PET imaging at 1, 24, 48 and 72 hr followed by ex vivo autoradiography, fluorescence microscopy and cryofluorescence tomography (CFT).

**Figure 4.**
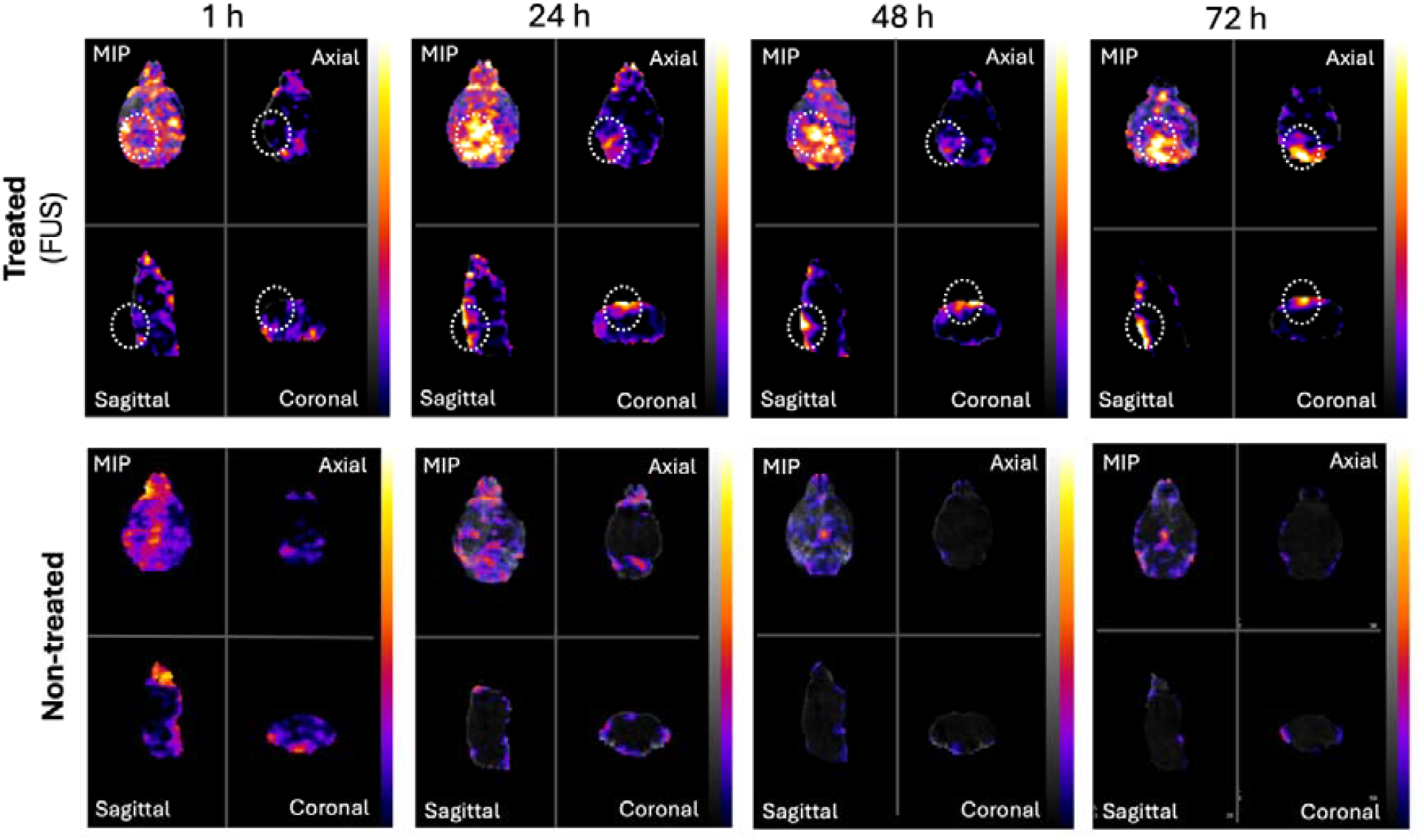
PET imaging of the brain showing delivery of [^89^Zr]Zr-Talidox to the brain post-treatment with focused ultrasound in the presence of microbubbles: FUS treated PET brain images with different views at t = 1 , 24 , 48 , 72 h (sonication target area circled in white), Bottom panel: Non-treated brain images.

Following confirmation that FUS-mediated BBB opening increases the delivery to the brain using PET imaging and biodistribution, we further determined the extent of delivery using PET image analysis and other imaging modalities, quantitatively and qualitatively.

### Quantification of brain opening and liposomal delivery to the brain

All FUS treatments were monitored in real time via passive cavitation detection. The detected energy increased when microbubbles entered the focal volume and decreased exponentially over time as a result of microbubble clearance (Figure 5A). Spectral analysis revealed consistent harmonic emissions throughout each treatment, indicating stable cavitation as well as sporadic broadband emissions indicative of inertial cavitation (Figure 5B-C). The mean energy from the acoustic emissions was 1.72 ± 1.11 × 10^-5^ V^2^s. The mean cavitation dose was 0.35 ± 0.09V, of which 89% came from stable cavitation and 11% inertial cavitation (Suppl. Fig. 1). Mean acoustic energy and total cavitation dose correlated linearly with whole brain uptake 72h post-treatment (R^2^ = 0.98 and 0.63 respectively, n = 4; Figure 4D and 4F). In addition, strong correlations between *in vivo* PET brain uptake and mean acoustic energy were found at 24h and 48h (R^2^ = 0.96 and 0.93, Figure 5E) and total cavitation dose at 48 h (R^2^ = 0.71, Figure 5G). The stable harmonic cavitation dose, stable ultraharmonic dose, and inertial cavitation doses did not correlate significantly with brain uptake (suppl. Fig. 1).

**Figure 5.**
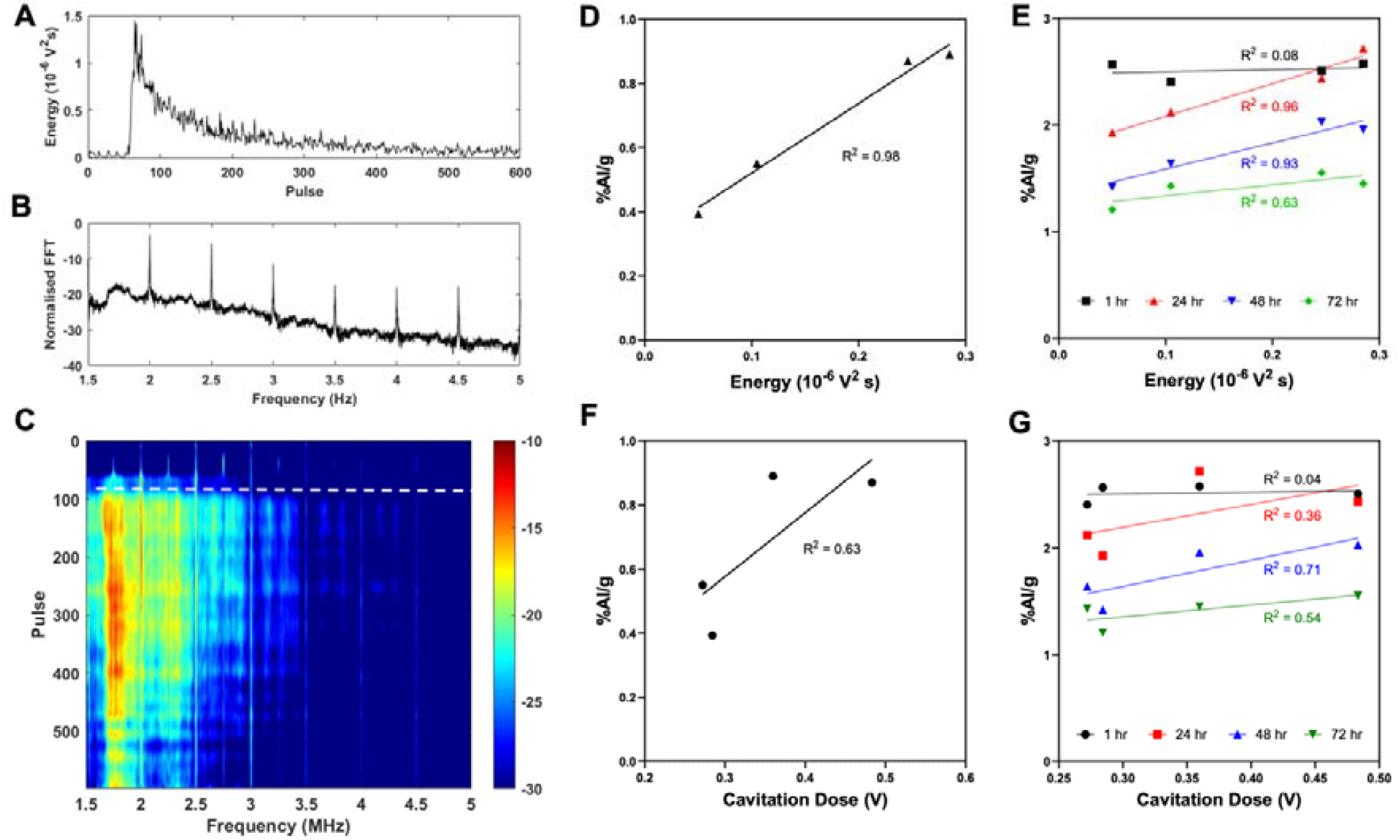
Mean acoustic energy and total cavitation dose correlate with liposome delivery 72 h post FUS: (A) Total energy per therapeutic pulse; (B) Normalised amplitude of FFT performed over cavitation emissions; (C) Spectrogram of FUS treatment (white dashed line indicates when microbubbles enter focal volume); (D, F) Linear correlation between mean acoustic energy and Cavitation dose with ex vivo brain uptake 72 h post treatment respectively; (E, G) linear correlations between mean acoustic energy and Cavitation dose with in vivo PET brain uptake.

The ex-vivo biodistribution of [^89^Zr]Zr-Talidox combined with the microbubble and FUS treatments was consistent in most organs with that described above, apart from the brain showing a 14-fold (14 ± 2) increase in the brain tissue accumulation of [^89^Zr]Zr-Talidox for the FUS-treated group (0.7 ± 0.2% IA/g) over the non-treated group (0.05 ± 0.02% IA/g) (Figure 6A,B). Using these quantitative biodistribution values, the doxorubicin drug dose concentration in the brain was determined as 0.8 ± 0.2 µg/g and 0.35 ± 0.1 µg as the absolute doxorubicin accumulation. The drug concentration accumulated in the brain after 72 h is 0.28 ± 0.09 % of the injected dose. This is an appreciable increase in the amount of cancer therapeutics that can be delivered to the brain without active targeting modification of liposomes and in cases of liposomal doxorubicin delivery using FUS.^42,67^

**Figure 6.**
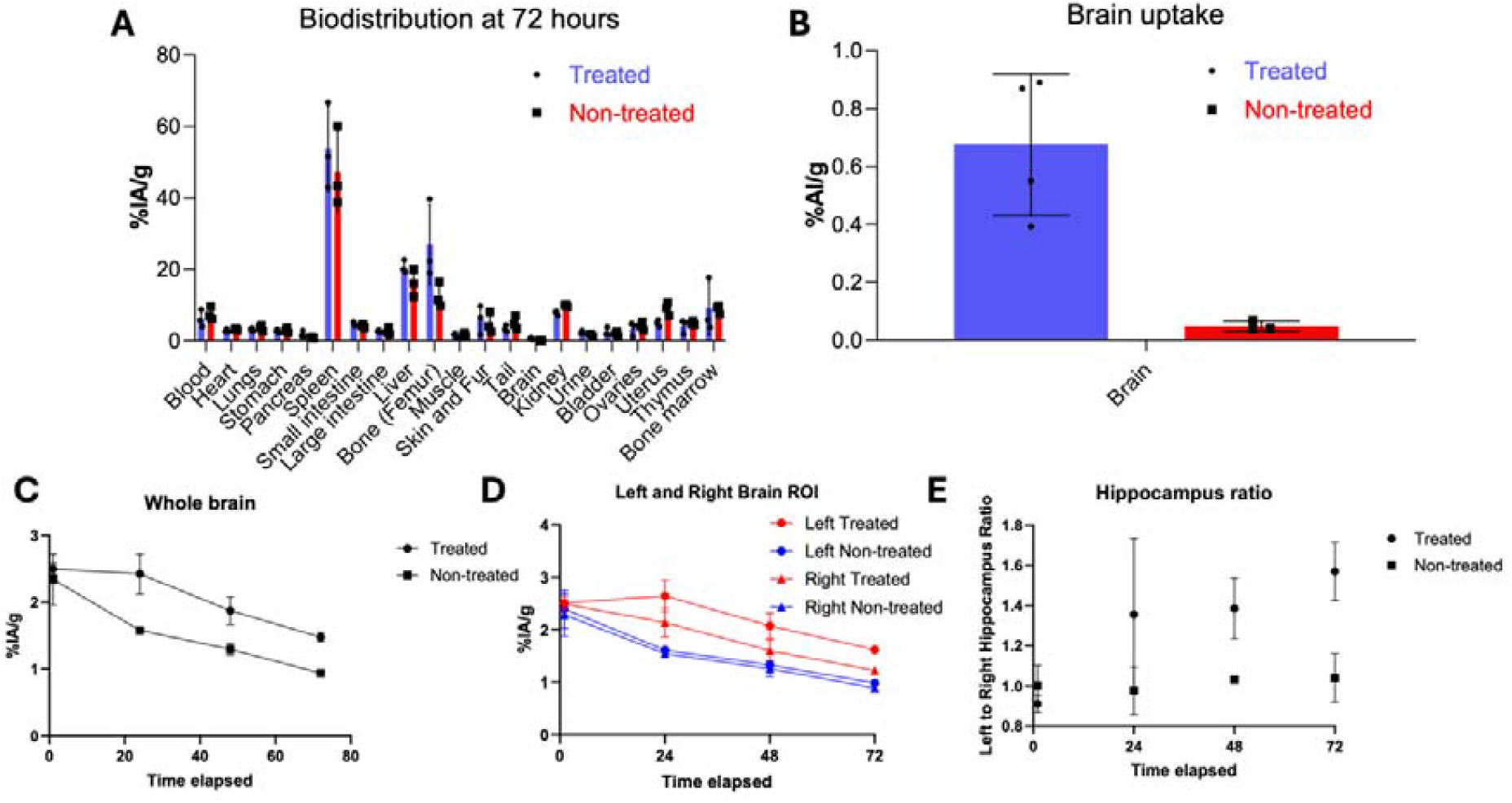
Biodistribution and image-based analysis comparison between FUS treated and non-treated brains: **(A)** Ex vivo biodistribution at t = 72 h comparing whole body biodistribution of [^89^Zr]Zr-Talidox between FUS treated and non-treated mice; **(B)** Ex vivo brain biodistribution at t = 72 h showing 14-fold increase in brain accumulation of Talidox in the treated brains compared to non-treated brains; **(C)** *In vivo* image-based quantification analysis of whole brain accumulation of [^89^Zr]Zr-Talidox over different time points at t= 1 hr, 24 hr, 48 hr, 72 hr; **(D)** *In vivo* image-based quantification analysis of [^89^Zr]Zr-Talidox accumulation observed in the left and the right hemisphere of treated and non-treated brains over different time points; (E) *In vivo* image-based quantification analysis of the ratio of uptake observed in the left hippocampus to the right hippocampus in treated and non-treated brains.

Further quantification using PET imaging allowed longitudinal tracking of the liposomal delivery in the brain and determination of regional biodistribution within different brain regions, including the comparison with the contralateral side. As seen in Figure 6C, differences in brain accumulation are evident throughout the different time points, showing the observed whole brain uptake was higher at each imaging time point. The observed uptake for the sonicated side of treated brains was higher than observed for the contralateral side after 24 h (2.5 ± 0.4 % IA/g vs 2.13 ± 0.1 % IA/g). Both sides of the treated brain showed higher uptake than untreated brains (Figure 6D). This confirmed that the BBB opening not only increased penetration of the liposomes into the brain parenchyma, but their smaller size allowed better distribution in the brain, thereby also showing increased uptake on the contralateral control side of the treated brains.

We evaluated the uptake in different regions of the brain, including the hippocampus, an area prone to neurological diseases, and its proximity to the zone of delivery using focused ultrasound. The left hippocampus of the sonication-treated brains showed long-lasting higher uptake ( 2.4 ± 0.1 % IA/g at t = 1 h, 2.28 ± 0.4 % IA/g at t = 24 h, 1.7 ± 0.4 % IA/g at t = 48 h, 1.33 ± 0.3 % IA/g at t = 72 h) compared to the left hippocampus of control untreated brains (2.5 ± 0.4 % IA/g at t = 1 h, 1.40 ± 0.02 % IA/g at t = 24 h, 1.10 ± 0.04 % IA/g at t = 48 h, 0.83 ± 0.1 % IA/g at t = 72 h). This further confirmed the targeted delivery of the liposomes and increased penetration towards the hippocampus. A comparison between the ratio of activity observed in left to right hippocampal regions between the treated and non-treated brains confirmed the targeted delivery enabled by focused ultrasound (Figure 6E). Further region-wise uptake of the different regions of treated and untreated brains can be found in the Supplementary information (Supp. Fig. 2).

To our knowledge, this is the first study exploiting PET imaging to non-invasively, longitudinally track and quantify the delivery of PET-labelled liposomes to the brain using FUS-mediated BBB opening. Previous studies have used optical imaging, MRI, or SPECT, which limit the quantification of the liposomal/drug dose delivered both in real time as well as post-mortem. ^31–38,41,42,45,46,68^ Differences in the extent of agent delivery have also been observed previously, depending on the amount of time these delivered agents are given to extravasate and diffuse within the brain. Other groups have investigated liposomal delivery at 0, 2, and 4 h after ultrasound exposure.^33,35,40,46^ Here, we explored liposomal distribution up to 72 h, with the maximum signal observed in the brain at 24 h. The decrease in signal at later time points suggest the BBB is no longer open, limiting further extravasation of liposomes into the brain. [^89^Zr]Zr-Talidox is also expected to be mostly cleared from the blood after 24 h. Moreover, these results could be explained by the reversible nature of the increase in BBB permeability using focused ultrasound.^69^

### Autoradiography, microscopy, and cryofluorescence tomography confirm delivery to the brain

Autoradiography is an ex-vivo imaging modality used to detect the biodistribution of radiotracers in tissue with high resolution (Figure 7). We performed the autoradiography on treated and untreated brains to determine the extent of delivery of [^89^Zr]Zr-Talidox®. The autoradiography images in Figure 7C confirmed the delivery of liposomes to the treated brains, while no delivery was observed for the control brains. Moreover, transversal sections (20 µm) of the whole brain (Suppl. Fig. 3) showed the delivery observed as multiple, homogenous, cloudy areas of radioactive signal within the targeted areas throughout the whole brain. No signal was observed in control brains injected with the same doses of [^89^Zr]Zr-Talidox liposomes and exposed to the phosphor plate for the same duration. This observation was consistent with previous studies that have shown similar patterns of liposome delivery using FUS.^33,35,36,40,46^ Multiple diffused delivery spots showing diffusion of the liposomes within the extracellular matrix following their extravasation from the blood vessels into the brain. This is likely due to the diffusion of these improved, smaller liposomes through the width of the extracellular matrix pores, which overcomes the issues faced by larger-sized conventional liposomal formulations^66^.

**Figure 7.**
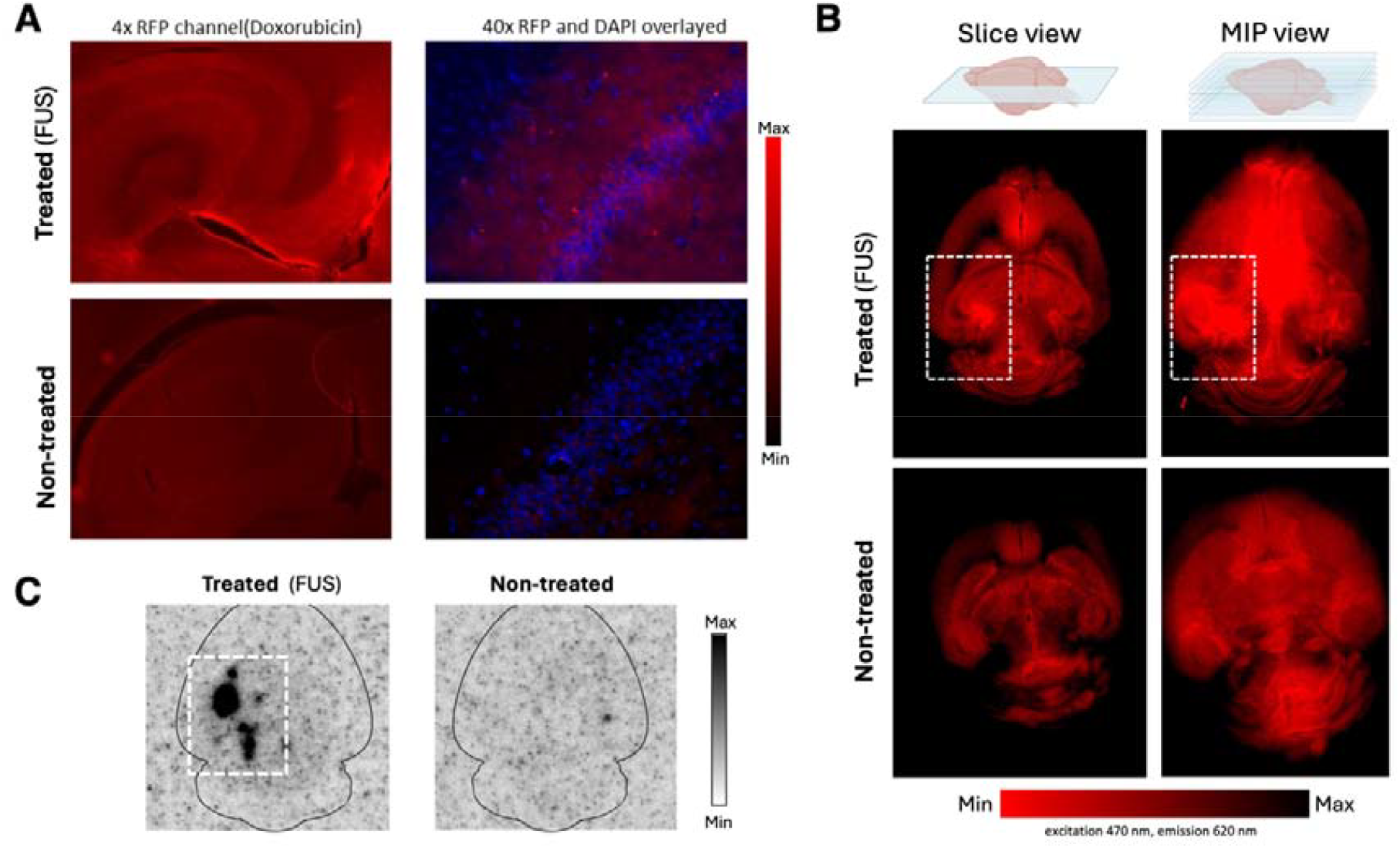
*Ex vivo* brain tissue imaging for visualisation of increased uptake of [^89^Zr]Zr-Talidox in FUS treated brains via: **(A)** fluorescence microscopy (doxorubicin fluorescence); **(B)** Whole-brain cryo-fluorescence tomography (CFT) (doxorubicin fluorescence), and **(C)** Autoradiography (^89^Zr)

Doxorubicin-loaded Talidox® liposomes exhibit fluorescence at RFP channel (Ex: 542/20 nm; Em: 593/40 nm). The microscopy images, as seen in Figure 7, showed enhanced fluorescent signal from the targeted regions (left hippocampus). The higher red fluorescence signal was also observed in regions of high cell concentration (DAPI signal). Thereby showing a higher accumulation of [^89^Zr]Zr-Talidox liposomes in regions with higher cell concentration. No uptake was observed in the untreated brain sections.

In addition to microscopy, which is limited to a single section and thereby limiting 3D visualization of the liposomal doxorubicin delivery within the brain, these limitations can be overcome using cryofluorescence tomography (CFT). An optical imaging modality, CFT, sequentially sectioned the tissue of interest, and each tissue section was imaged block-face using a fluorescence microscope. In this way, the 3D biodistribution of the object of interest (in our case Doxorubicin loaded on Talidox®) can be captured across the entire tissue. In the maximum intensity projection (MIP) of sonication-treated brains, bright fluorescence signal was observed on the sonicated left side of the brain, whereas background fluorescence was obtained for the non-treated contralateral right side. The background autofluorescence of the brain tissue limited our ability to quantify the delivered dose of doxorubicin from the acquired fluorescence. Moreover, the untreated brain with and without the administration of the [^89^Zr]Zr-Talidox did not show any increased signal in the brain. There was no relative difference between the observed fluorescence signal from the two brain hemispheres.

However, this study has potential limitations. The unlimited depth of detection and quantitative nature of PET is achieved at the cost of administering ionizing radiation and limited spatial resolution (preclinical (0.5-1 mm), clinical (4-5 mm)). The passive cavitation detection in our study was spatially limited to 1-D and provided lower resolution than 2-D/3-D passive acoustic mapping. For animal experiments, the number of animals was low for observing correlation with acoustic data.

In conclusion, this study introduced PET imaging for performing quantitative, non-invasive, longitudinal tracking of liposomes to the brain following focused ultrasound-mediated opening of the BBB. It not only allowed the determination of the extent of delivery over time but also the precise quantification of cancer therapeutic delivered to the targeted region of the brain. This study also establishes the safety of doxorubicin dose delivered to the brain using focused ultrasound. Future work will focus on investigating whether therapeutic concentrations of Doxorubicin delivered by these smaller, more stable, and longer circulating novel Talidox® liposomes exhibit therapeutic effects in tumor models of glioblastoma with an intact BBB. Further effects of increased dosing, repeating dosing, and longitudinal FUS treatments will also be investigated in aggressive models of tumors with lower survival, such as models of pediatric tumors. In addition, the effect of microbubbles on spleen blocking to aid the increased delivery of liposomal nanomedicines to sites of tumor and inflammation will also be investigated, as this can open an effective method to evade the reticuloendothelial system. Finally, further optimization of the cryoFT protocol will allow for quantitative assessment of liposomal accumulation in the brain.

## Conclusions

This study introduces simple radiolabelling of clinically relevant liposomes, facilitating the use of PET imaging as a non-invasive, quantitative method to longitudinally track the delivery of a novel, smaller liposomal formulation Talidox® to the brain via FUS-mediated BBB opening. PET imaging of [^89^Zr]Zr-Talidox and biodistribution analysis revealed that FUS effectively increased BBB permeability, significantly improving Talidox® delivery to the brain by 14-fold. The increased accumulation observed in PET imaging correlated with passively monitored cavitation dose and energy. Regional analysis showed effective liposome distribution preferentially on the treated side of the brain, especially in critical areas like the brain’s memory centre hippocampus.

Additional imaging techniques, such as autoradiography, microscopy, and cryo-fluorescence tomography, further validated the delivery of radiolabelled liposomes and the enclosed doxorubicin within the brain. Higher fluorescence was observed on the treated side within cell-dense regions due to increased delivery of doxorubicin in these regions. The administration of microbubbles did not interfere with overall liposome biodistribution, though a decreased splenic uptake was observed, which can be exploited further for applications in drug delivery.

Overall, this study highlights the potential of small Talidox® liposomes for efficient brain delivery of clinically relevant doses of doxorubicin using FUS, with delivery monitored longitudinally and quantitatively using PET imaging. This study will lay the groundwork for further research on therapeutic effects in brain cancer and optimization of delivery strategies to enhance treatment outcomes.

## Acknowledgments

This work was supported by the Centre of Excellence in Medical Engineering, funded by the Wellcome Trust and the Engineering and Physical Sciences Research Council (EPSRC) (grant number WT 203148/Z/16/Z); EPSRC Programme Grant (EP/S032789/1 ‘‘MITHRAS’’), Children’s Cancer and Leukaemia Group (CCLG)/Little Princess Trust (CCLGA 2022 25), Action Medical Research/LifeArc (GN3017), Focused Ultrasound Foundation (FUS1050R1), and Abbie’s Army/Children’s Brain Tumor Drug Delivery Consortium (KCL/G12/22). The authors acknowledge the support of InnoMedica Holding AG for providing Talidox liposomes used in this study. PET scanning equipment at KCL was funded by an equipment grant from the Wellcome Trust under grant no. WT 084052/Z/07/Z. Radioanalytical equipment was funded by a Wellcome Trust Multiuser Equipment grant: a multiuser radioanalytical facility for molecular imaging and radionuclide therapy research. The authors acknowledge support from the National Institute for Health Research (NIHR) Biomedical Research Centre based at Guy’s and St Thomas’ NHS Foundation Trust and KCL (grant no IS-BRC-1215-20006). The authors would also like to thank Dr. Victor Jeannot from Bracco SpA for supplying the BR1 microbubbles used in this project, as well as Dr. Kavitha Sunassee and Dr. Adam Jones for their help and support with the preclinical facility. The authors would finally like to thank Dr. Peter Gawne (Barts Cancer Institute, Queen Mary University of London) and Dr. Filipa Mota (Perspective Inc.) for facilitating access and running samples on the Cryo Fluorescence Tomography (CFT) equipment based at Barts Cancer Institute, QMUL. All graphical figures have been created using Biorender.com with a publication license.

## GRAPHICAL ABSTRACT

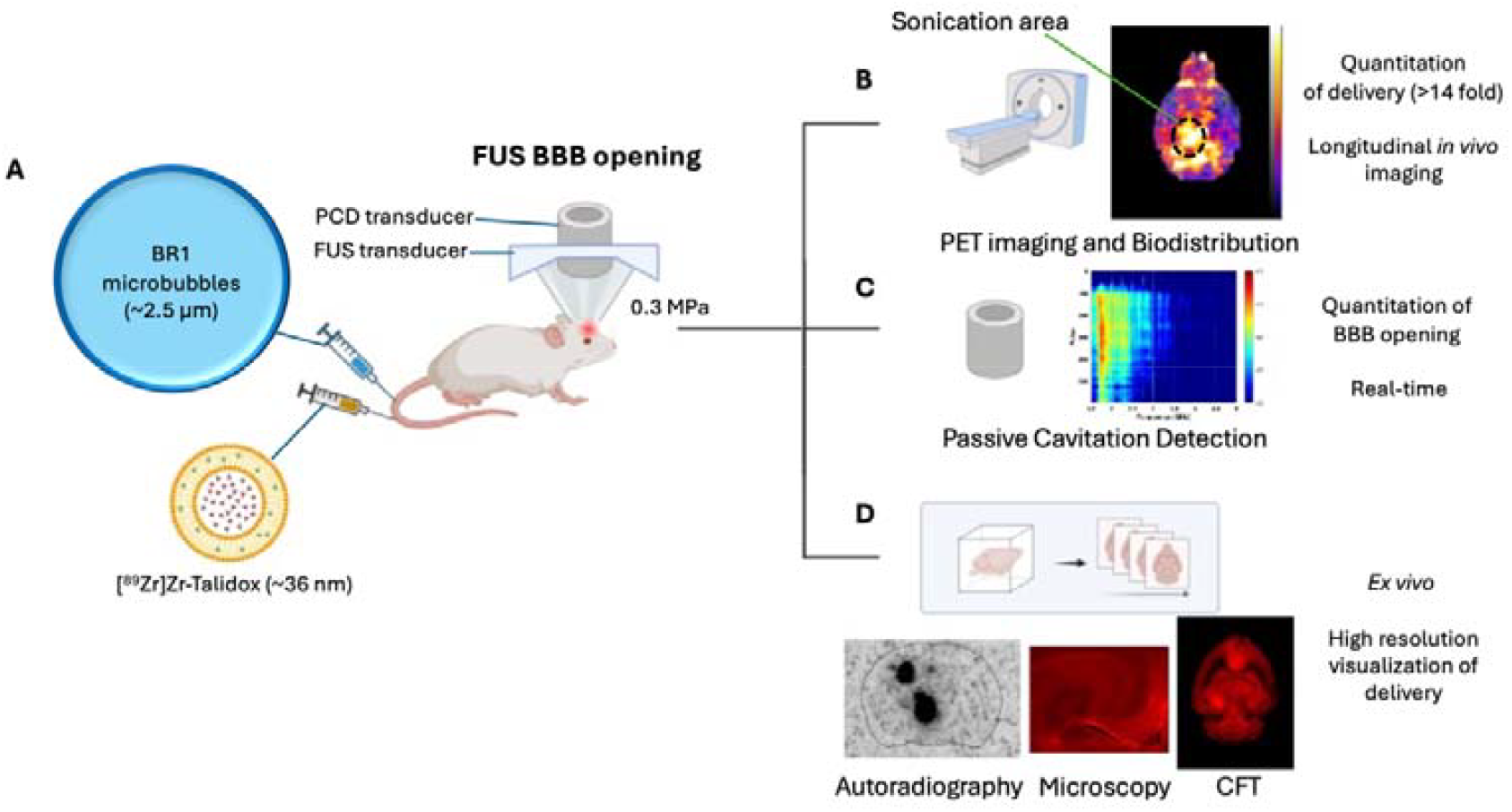

