## Supplemental Information for "PET imaging for non-invasive monitoring of ^89^Zr-Talidox delivery to the brain following focused ultrasound-mediated blood-brain barrier opening"

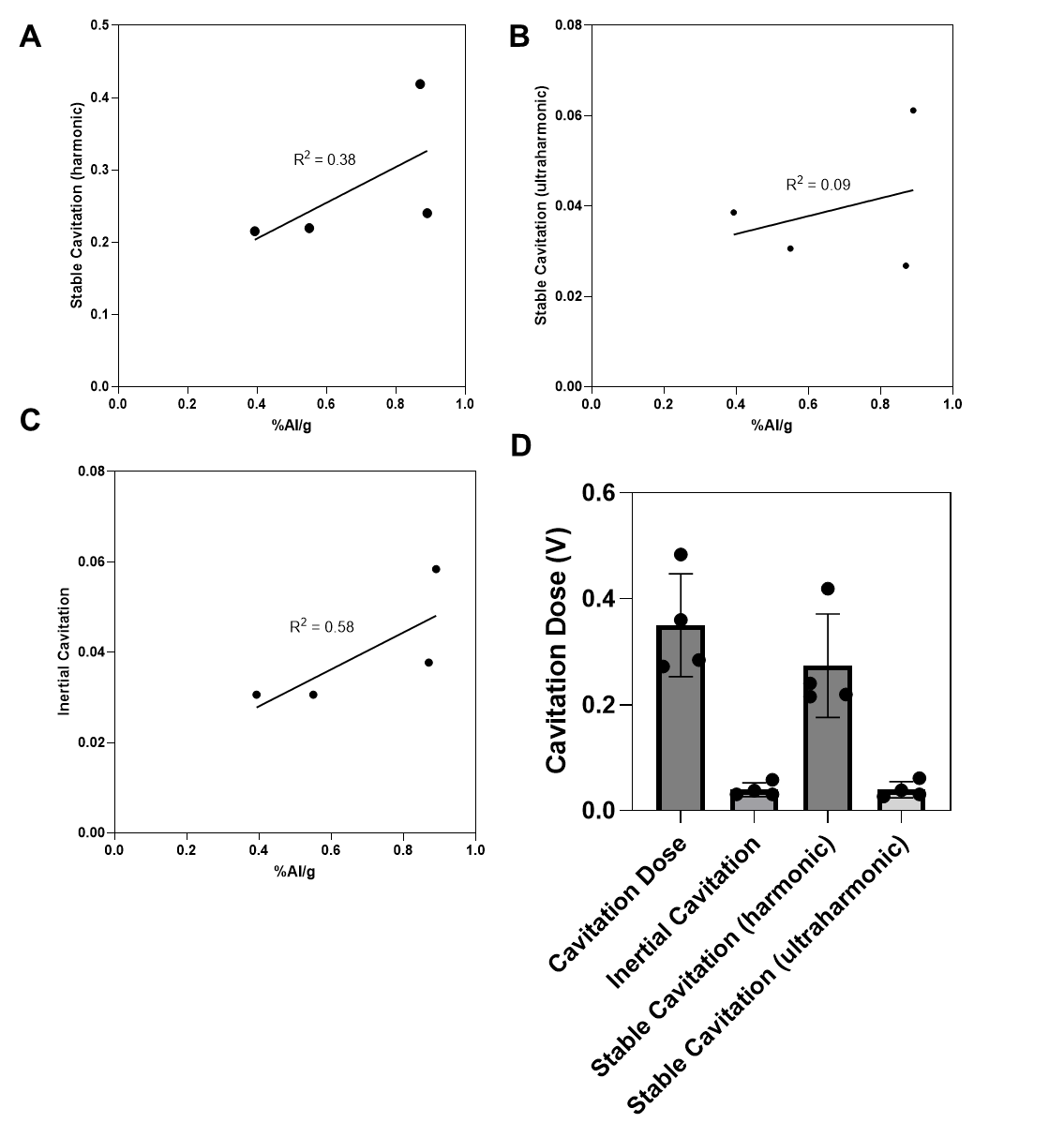

### Correlation between cavitation dose and Talidox dose delivered

***Figure S1.*** ***(A-C)*** *linear correlation of stable harmonic, stable ultraharmonic and inertial cavitation dose;* ***(D)*** *Mean total Cavitation dose, inertial cavitation, stable harmonic, stable ultraharmonic dose.*

### Brain PET imaging analysis

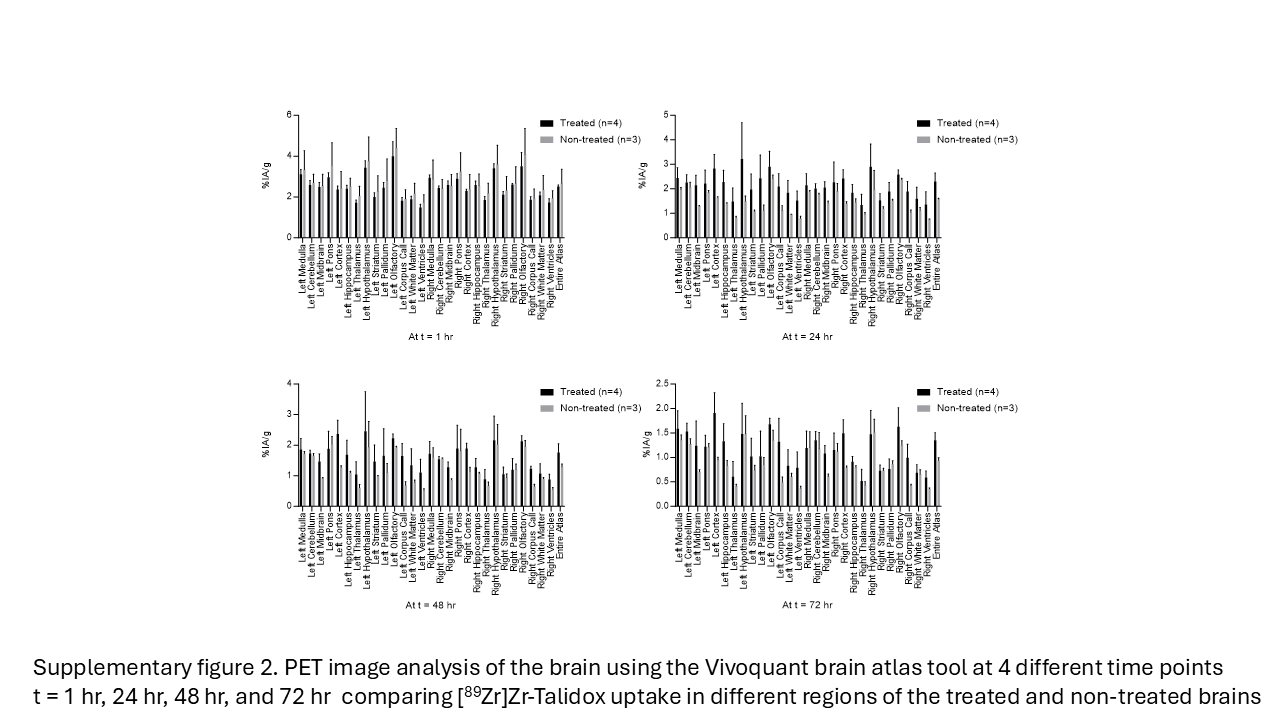

**Figure S2.** PET image analysis of the brain using the VivoQuant brain atlas tool at 4 different time points at t = 1 h, 24 h, 48 h, and 72 h comparing [^89^Zr]Zr-Talidox uptake in different regions of the treated and non-treated brains.

### Qualitative visualisation of radioactive signal in the brain using autoradiography

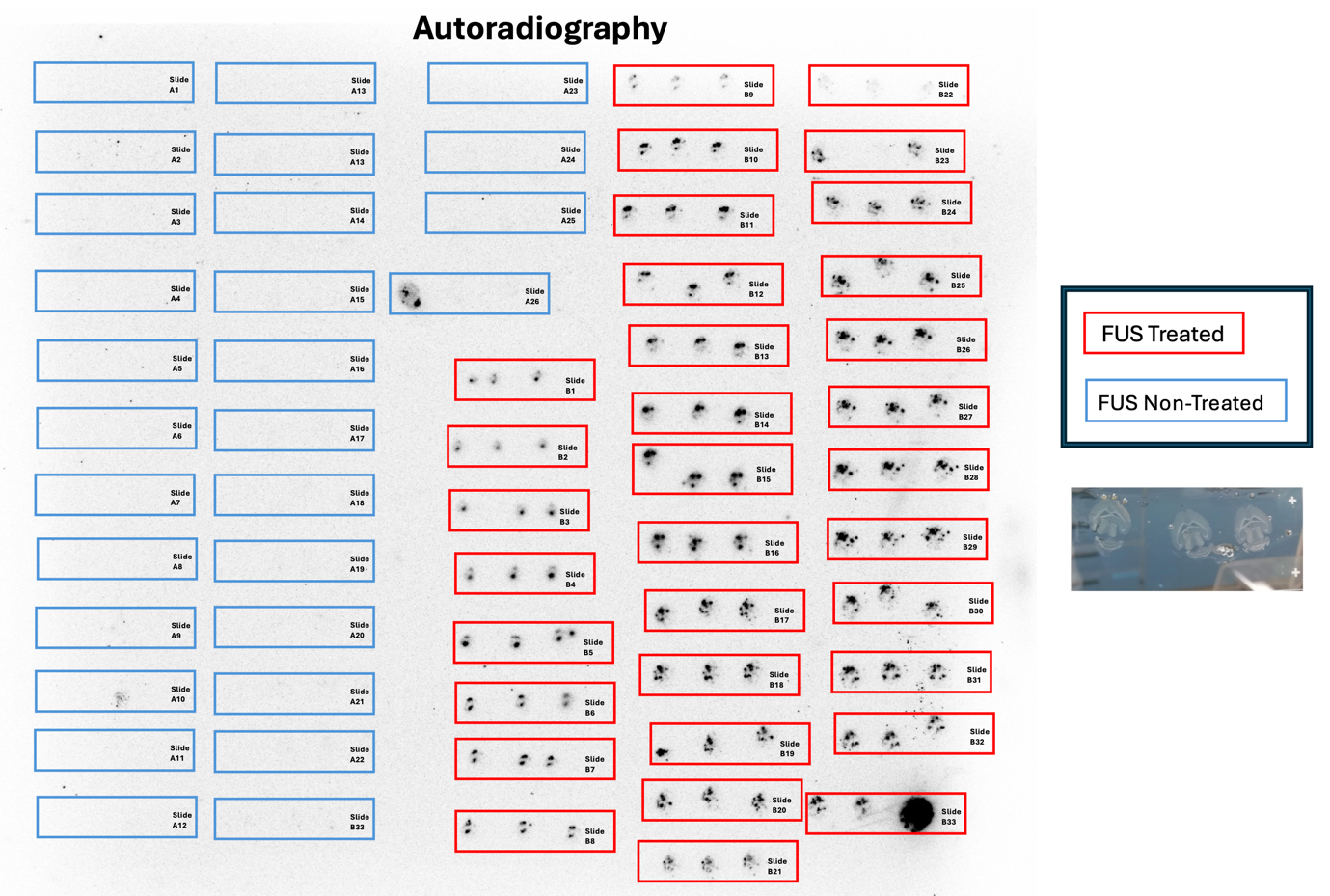

Figure S3. **Autoradiography of focused ultrasound treated and untreated brains showing comparative localization of [^89^Zr]Zr-Talidox within the brain**: Slides A1-A26 (blue) show autoradiography of 20 µm brain sections x 3 per slide of untreated brains with increasing numbers depicting the increasing depth from top to bottom of the axial plane; Slides B1-B33 (red) show autoradiography of 20 µm brain sections x 3 per slide of treated brains with increasing numbers depicting the increasing depth from top to bottom of the axial plane. A visible light representative image showing brain sections placed on a glass slide.
